# Key contribution of prefrontal inhibition to passive coping behaviour: chronic stress and fast-acting antidepressant

**DOI:** 10.1101/2024.10.20.619255

**Authors:** Tsz Hei Fong, Tianxiang Li, Xiaoyan Ma, Xiang Cai, Qiang Zhou

## Abstract

Persistent passive coping behaviour is a hallmark feature in major depression and is reversed by fast-acting antidepressants (such as ketamine). This behaviour is regulated by a specific cortico-midbrain circuit. However, whether the prefrontal cortex (PFC), especially inhibition in PFC, contributes to the modulation of passive coping, and whether this modulation is important for mediating the impacts of chronic stress and/or fast-acting antidepressants, are poorly understood. Here, we found that rostral prelimbic cortex (rPL) bidirectionally controls the occurrence of passive coping behaviour where excitatory and inhibitory neurons play opposite roles. Chronic stress leads to reduced excitation/inhibition (E/I) ratio, reflected as alterations in *in vivo* spiking rate, synaptic inputs and intrinsic excitability of both excitatory and inhibitory neurons. A fast-acting antidepressant, (2*R*, 6*R*)-hydroxynorketamine (HNK), reduced passive coping behaviour, restored rPL E/I ratio and partially reversed altered properties in rPL neurons, in chronically stressed mice. Importantly, chronic stress and HNK mostly affected fast-spiking/parvalbumin inhibitory neurons instead of other inhibitory neurons, indicating the important role of this subtype of inhibitory neurons in the above processes. These findings demonstrate the importance of rPL E/I balance in regulating passive coping, which can be modulated by chronic stress and rapidly restored by fast-acting antidepressant.

## INTRODUCTION

Chronic stress is one of the most relevant precipitating risk factors for mental illness [1]. Adequate adjustments at psychological, physiological, and behavioural levels are required to maintain homeostatic stability and to avoid genesis of disorders [2]. Appraisal of and adaptation to stress are critical to human and animal survival. Chronic maladaptation to stress may lead to a range of stress-related disorders, including major depression, post-traumatic stress disorder and anxiety, which have severe impacts on life quality and even the life span of the affected [3, 4]. One widely used parameter to measure this maladaptation to stress is passive coping (p-coping) in face of stress stimuli. P-coping is defined as a low-effort, avoidant, or submissive response to stressors, which includes withdrawal, avoidance, denial, or disengagement [5]. Since improvement in p-coping behaviour is widely used in the dissection of the related processes/mechanisms and in evaluating the efficacy of treatment of the stress-related disorders (especially depression), a better understanding of the circuity and key target neurons modulating p-coping behaviour is critical. This is one main aim of the current study.

Prefrontal activity has an important contribution to the regulation of stress responses and p-coping behaviour, especially via rostral prelimbic (rPL) output glutamatergic neurons [6, 7]. The rPL projections to anteroventral bed nuclei of the stria terminalis (avBNST) and ventrolateral periaqueductal gray (vlPAG) selectively modulate the emergence of passive coping behaviour [8, 9]. To achieve fine-tuning of behaviours, inhibitory neurons in the prefrontal cortex (PFC) coordinate local neuronal networks and gate the outputs of excitatory neuron activity under both adaptive and pathological conditions [10]. However, whether inhibitory neurons in rPL may play such a role in p-coping is not well understood. Chronic stress resulted in enhanced passive coping behaviour [11], together with a significant increase in the activity of inhibitory neurons and pre-synaptic GABA release in PFC [12, 13], but whether these two events are casually related is unclear. We have addressed the above two questions in the current study.

The fast-acting antidepressants, represented by ketamine, have shown a rapid relief from treatment-resistant major depression in human patients [14, 15] and a rapid reversal of depression-like behaviours including p-coping behaviour in rodent depression models [16]. For example, Chen, et al. [17] reported decreased PL-avBNST excitatory activities and increased immobility time in FST and TST, in a chronic restrained mice model, and all these alterations were reversed by ketamine treatment. Due to the significant side effects of ketamine (dissociation, and potential toxicity associated with long-term use) [18], a search for compounds that preserve ketamine’s anti-depressant efficacy but lack ketamine’s side effects is in hot pursuit. A promising candidate is (2*R*, 6*R*)-hydroxynorketamine (HNK), a major metabolite of ketamine, that has shown anti-depressant efficacy comparable to ketamine in rodent depression models [19–21] and satisfactory Phase I trial results [22]. But whether HNK may reverse several or all of the chronic stress-induced alterations in the rPL circuitry related to p-coping is not understood. We have addressed this question.

To answer the above questions, we have used a combination of methods, from *in vivo* and *in vitro* recordings of neuronal activity and properties, to opto- and chemo-genetic manipulations, and behavioural measurements. In rodents, immobility under inescapable stressful circumstances, such as forced swimming test (FST) and tail suspension test (TST), represents the p-coping behaviour.

In this study, we used a treatment-resistant mice model of depression via a 14-day injection of adrenocorticotrophic hormone (ACTH). We report that rPL inhibitory neurons, especially PV-neurons, bidirectionally modulate the emergence and duration of p-coping (TST). Chronic stress alters the spike rate and properties of both rPL excitatory and inhibitory neurons, and part of these alterations is rapidly reversed by HNK treatment which is associated with restored p-coping.

## RESULTS

### Chronic stress increases passive coping and alters rPL excitation and inhibition

We used a well-established chronic stress model via intraperitoneal (i.p.) injection of ACTH for 14 days (Fig. 1A) in BALB/c mice [23], which showed enhanced depressive-like behaviours including TST, FST and SPT (Fig. 1B and S-Fig. 1A). The FST and TST are considered as measures of p-coping behaviour [6, 7]. Since TST does not require mice to be immersed in water which presents a technical challenge for recording and imaging *in vivo*, we have selected it as our main model of testing p-coping. We found the increased TST immobility time is caused by longer duration of each immobile episode rather than by the increased number of immobile episodes (Fig. 1C).

**Figure 1.**
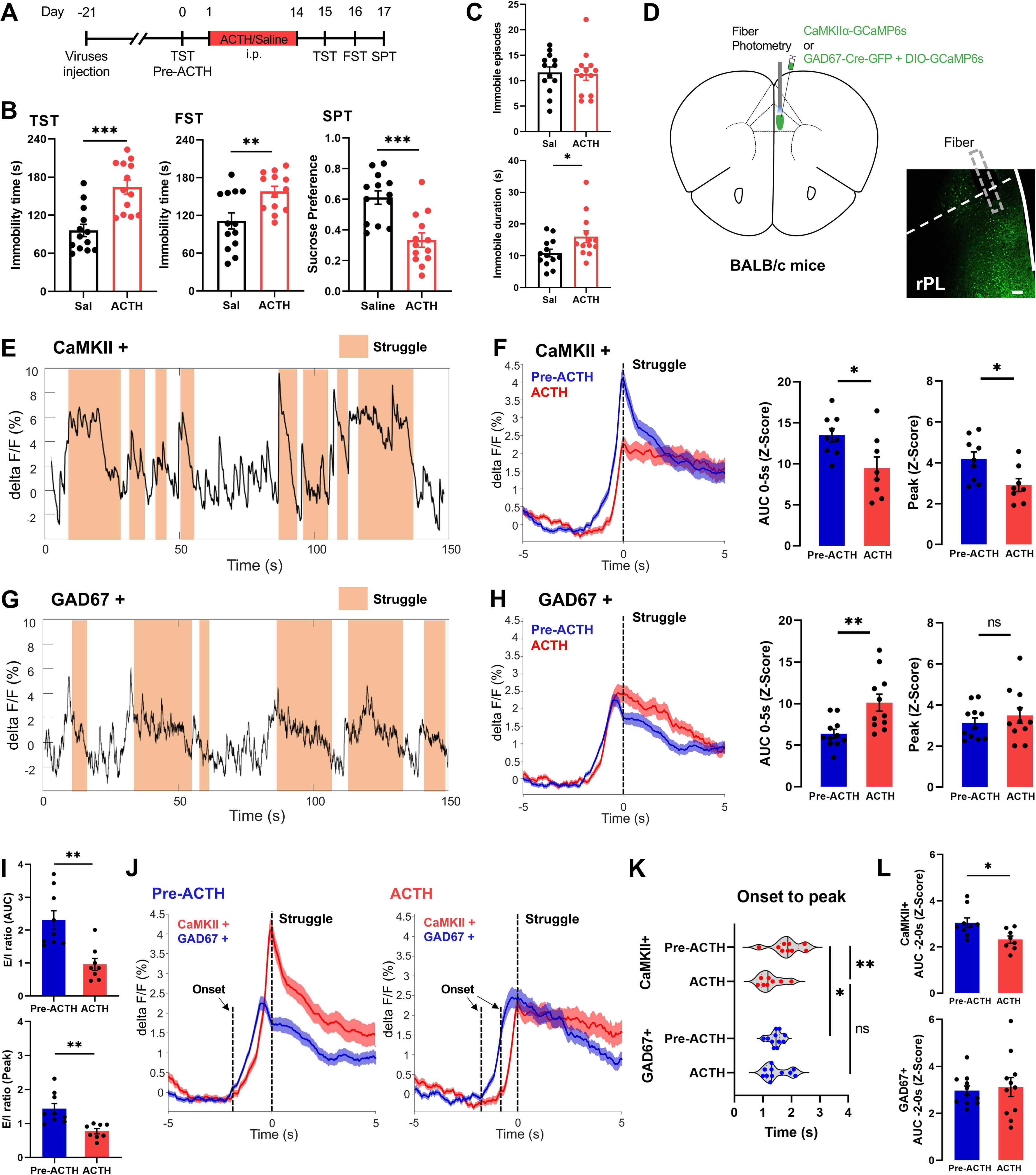
Chronic stress induces depression-like behaviours and alters rPL activity. (A) Experiment protocol for model establishment and behavioural tests. (B) Chronic ACTH injection induced alterations in TST, FST and SPT (Saline [Sal, *n* = 13 mice] vs. ACTH [*n* = 13]). (C) Numbers of immobility episodes (upper) and averaged duration of each immobility episode (bottom) during TST in saline- and ACTH-injected mice. (D) Location of virus injection and sample image of GCaMP6s injection/optical fiber position (gray dotted rectangle) in rPL of BALB/c mice. Scale bar, 100 μm. (E, G) Representative Ca^2+^ responses in CaMKII+ (E) or GAD67+ (G) neurons during TST, with struggling periods indicated by shaded regions. (F, H) (Left) Averaged Ca^2+^ responses (mean ± SEM) in CaMKII+ [F, *n* = 9 mice] or GAD67+ [H, *n* = 11] neurons before and after ACTH injection in the same set of mice. Traces were aligned to the onset of struggling (time 0). (Right) Area under curve (AUC) from 0 to 5 s and peak values of Ca^2+^ responses (pre-ACTH vs. ACTH). (I) E/I ratio calculated from Ca^2+^ responses in by dividing AUC (upper) or peak values (bottom) of CaMKII+ neurons to that in GAD67+ neurons (pre-ACTH vs. ACTH). (J) Averaged Ca^2+^ responses (mean ± SEM) in CaMKII+ and GAD67+ neurons before (left) and after (right) ACTH injection. (K) Quantification of durations in Ca^2+^ responses, from response onset to peak, in CaMKII+ or GAD67+ neurons before and after ACTH injection. (L) AUC (from −2 to 0 s) of Ca^2+^ responses in CaMKII+ (upper) or GAD67+ (bottom) neurons (pre-ACTH vs. ACTH). Data are shown as mean ± SEM. Unpaired *t*-test (B, C, F, H, I, K and L). *, *p* < 0.05; **, *p* < 0.01; ***, *p* < 0.001; ****, *p* < 0.0001; ns, not significant.

Previous studies have shown that rPL plays an important role in p-coping [8], we first monitored rPL activity in ACTH mice by measuring the Ca^2+^ responses in rPL excitatory and inhibitory neurons during TST, via expression of GCaMP6s in these neurons (Fig. 1A, D and S-Fig. 1B). Photometry recording revealed that both excitatory (CaMKII+) and inhibitory (GAD67+) neurons showed time-locked responses to the onset of struggling (Fig. 1E-H and S-Fig. 1C). We observed significantly smaller peak and area under the curve (AUC) in Ca^2+^ responses in the CaMKII+ neurons after ACTH injection (Fig. 1F), and significantly larger AUC in GAD67+ neurons (Fig. 1H), suggesting opposite changes in their responses as a result of chronic stress. We then divided AUC and peak values of CaMKII+ neurons over that of GAD67+ neurons to calculate an E/I ratio, and we observed a significant reduction (Fig. 1I). This reduced E/I ratio may contribute to the increased duration of immobility episodes.

The PFC inhibitory neurons connect to their neighboring excitatory neurons to fine-tune behaviour [24]. When Ca^2+^ responses in the CaMKII+ and GAD67+ neurons were compared before and after ACTH injection (Fig. 1J), we observed that the peak of Ca^2+^ response in CaMKII+ neurons time-locked to the struggling onset following an earlier peak of GAD67+ neurons (S-Fig. 1D). We found that Ca^2+^ response onset was significantly delayed in CaMKII+ neurons in the ACTH-injected mice, resulted in a shorter onset to peak and reduced AUC (−2 to 0 s) in CaMKII+ neurons (S-Fig. 1E and Fig. 1K-L). However, onset time, onset to peak and AUC in GAD67+ neurons were not altered in ACTH mice (S-Fig. 1E and Fig. 1K-L), leading to a higher inhibition before struggling onset. These results suggest that excitation before struggling influences the p-coping duration.

### Chronic stress affects both rPL excitatory and inhibitory neurons

The above results indicate altered responses in rPL excitatory and inhibitory neurons during TST in chronically stressed mice. To examine whether the activity and/or properties of these neurons are altered by chronic stress, we first performed *in vivo* electrophysiological recordings from the same set of neurons before and after ACTH injection, using previously established criteria to distinguish between excitatory and inhibitory neurons (S-Fig. 2A-B) [25]. After ACTH injection, a significantly higher spike rate was observed in the inhibitory neurons starting from day 6 – 9 while a significantly lower spike rate in the excitatory neurons starting from day 12 – 14 (Fig. 2A). These opposing changes led to a reduced E/I ratio starting from day 6 – 9 (Fig. 2B). Notice that alterations in the inhibitory neurons occur prior to that in the excitatory neurons.

**Figure 2.**
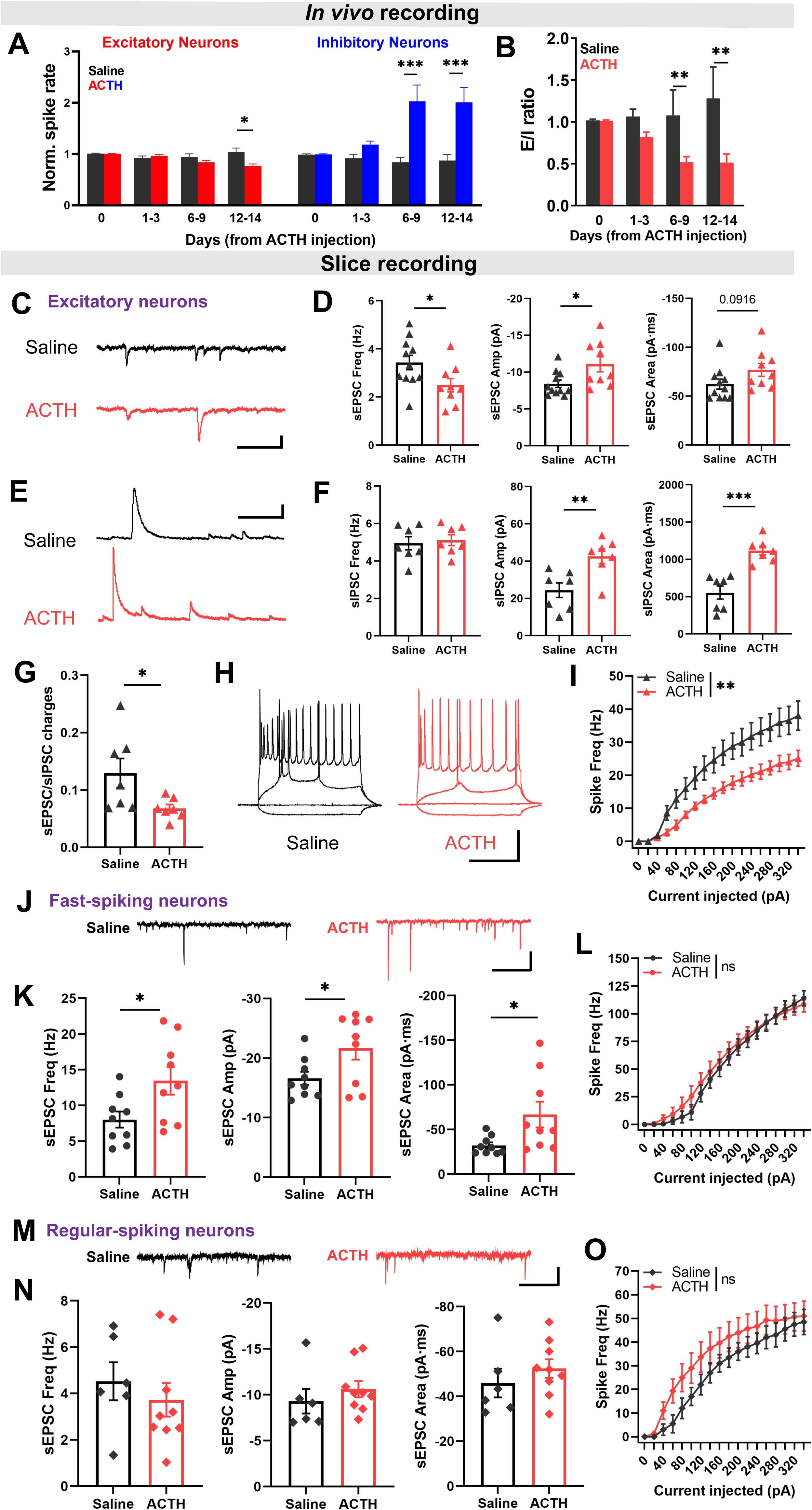
Chronic stress alters the properties of rPL excitatory and inhibitory neurons and E/I balance. (A) Changes in *in vivo* spike rate in excitatory [*n* = 100 - 150 cells/group] and inhibitory [*n* = 30 - 60 cells/group] neurons from saline- [*n* = 5 mice] or ACTH-injected mice [*n* = 6 mice], normalized to spike rates on day 0 (Saline vs. ACTH). (B) E/I ratio (spike rate of excitatory over inhibitory neurons) calculated from (A). (C, D) Representative sEPSC traces (C) and summary data (D) for sEPSC frequency, amplitude, and area in excitatory neurons from saline- [*n* = 11 neurons/3 mice] or ACTH-injected mice [*n* = 9 neurons/3 mice] (Saline vs. ACTH). Scale bars, 10 pA and 250 ms. (E, F) Representative sIPSC traces (E) and summary data (F) for sIPSC frequency, amplitude and area in excitatory neurons from saline- [*n* = 7 cells/3 mice] or ACTH-injected mice [*n* = 7 cells/3 mice] (Saline vs. ACTH). Scale bars, 40 pA and 250 ms. (G) E/I ratio of synaptic responses calculated by dividing total areas of sEPSC over sIPSC in the same excitatory neurons from saline- [*n* = 7 cells/3 mice] or ACTH-injected [*n* = 7 cells/3 mice] mice. (H) Representative traces of action potentials elicited by current injections in excitatory neurons (Black: Saline; Red: ACTH). Current steps, −40, 0, 40, and 340 pA. Scale bars, 30 mV and 250 ms. (I) Intrinsic excitability (injected currents −40 to 340 pA) in excitatory neurons from saline- [*n* = 11 cells/3 mice] or ACTH-injected mice [*n* = 10 cells/3 mice]. (J, K) Representative sEPSC traces (J) and summary data (K) for sEPSC frequency, amplitude and area in fast-spiking GAD67+ neurons from saline- [*n* = 9 cells/3 mice] or ACTH-injected mice [*n* = 9 cells/3 mice]. Scale bars, 30 pA and 250 ms. (L) Intrinsic excitability in fast-spiking GAD67+ neurons from saline- [*n* = 9 cells/3 mice] or ACTH-injected mice [*n* = 9 cells/3 mice]. (M, N) Representative sEPSC traces (M) and summary data (N) for sEPSC frequency, amplitude, and area in regular-spiking GAD67+ neurons from saline- [*n* = 6 cells/3 mice] or ACTH-injected mice [*n* = 9 cells/4 mice]. Scale bars, 10 pA and 250 ms. (O) Intrinsic excitability in regular-spiking GAD67+ neurons from saline- [*n* = 6 cells/3 mice] or ACTH-injected mice [*n* = 9 cells/4 mice]. Two-way ANOVA (A, B, I, L and O). Unpaired *t*-test (D, F, G, K and N).

The above-observed alterations in neuronal spiking are caused by altered synaptic inputs and/or neuronal intrinsic excitability. To examine this, we recorded from neurons that selectively participated in the TST by using the Tet-Off system with injection of cFos-tTA and TRE-mCherry virus in rPL (S-Fig. 2C). We observed a significantly lower spontaneous excitatory post-synaptic current (sEPSC) frequency, larger sEPSC amplitude with no alteration in sEPSC area and decay time (Fig. 2C-D and S-Fig. 2D), and a significantly higher inhibitory post-synaptic current (sIPSC) amplitude and area (Fig. 2E-F) in excitatory neurons from ACTH mice. These results are in general agreement with the lower E/I ratio at the synaptic level in the excitatory neurons (Fig. 2G). In addition, the intrinsic excitability, measured using depolarizing current injections, was significantly lower in excitatory neurons from ACTH mice (Fig. 2H-I). Both alterations indicate reduced outputs from these excitatory neurons.

Next, we recorded from rPL inhibitory neurons. We initially attempted to record from neurons expressing both c-Fos and GAD67, but there were very few such neurons in the superficial layer of PFC. Thus, we recorded randomly from GAD67+ neurons which can be separated into fast-spiking neurons (FSN) and regular-spiking neurons (RSN). In FSNs from ACTH mice (highest spike rate > 60 Hz), we observed a significantly higher sEPSC frequency, amplitude, and area (Fig. 2J-K), with no significant change in sEPSC decay time (S-Fig. 2E) or intrinsic excitability (Fig. 2L). In addition, we did not find any significant changes in any of the parameters we measured in RSNs from the same mice (Fig. 2M-O and S-Fig. 2F). Taken together, FSNs appear to be the main inhibitory neuron population that is significantly affected by chronic stress.

Although we cannot record from c-Fos+ and parvalbumin (PV)+ neurons, we can use staining in the brain sections to determine whether the density of PV-neurons participating in the TST is altered by chronic stress. We performed c-Fos labelling in GAD67-GFP mice (S-Fig. 2C). Since about 90% of rPL FSNs also express parvalbumin [26], we used PV-selective antibodies to identify them (S-Fig. 3A). ACTH did not alter the density of PV+ or c-Fos+ neurons in rPL, but significantly increased the density of PV+ neurons that were co-labeled with c-Fos (S-Fig. 3B-D). Put together, chronic stress affects both excitatory and inhibitory neurons in the rPL and decreases rPL E/I ratio, promoting rPL inhibition via increased participation of PV-neurons during TST which may mediate the more prominent p-coping behaviour observed in the mice.

### The rPL inhibition promotes passive coping behaviour

We have observed alterations in p-coping behaviour and altered activity and properties in both rPL excitatory and inhibitory neurons in chronically stressed mice. Are these alterations connected? Put another way, does rPL neuron activity modulate p-coping behaviour? We expressed excitatory channel-rhodopsin-2 (ChR2) or inhibitory halorhodopsin (NpHR) in CaMKII+ and mDlx+ neurons to manipulate their activity level using light stimulation during TST (Fig. 3A-B, E). We confirmed the effectiveness of the viral expression using PFC slices (S-Fig. 4A, C, E, G). Pulsed (20 Hz) blue light stimulation of excitatory neurons reduced total immobility time in TST in both saline- and ACTH-injected mice, compared to the same mice without light stimulation (Fig. 3C). For the immobility time during all light OFF or ON periods, immobility time during light ON periods was significantly lower (S-Fig. 4B), suggesting that rPL excitation instantly promotes struggling and suppresses p-coping behaviour. Yellow light stimulation (step) of rPL excitatory neurons increased total immobility time without causing any rapid change in p-coping behaviour (Fig. 3D and S-Fig. 4D).

**Figure 3.**
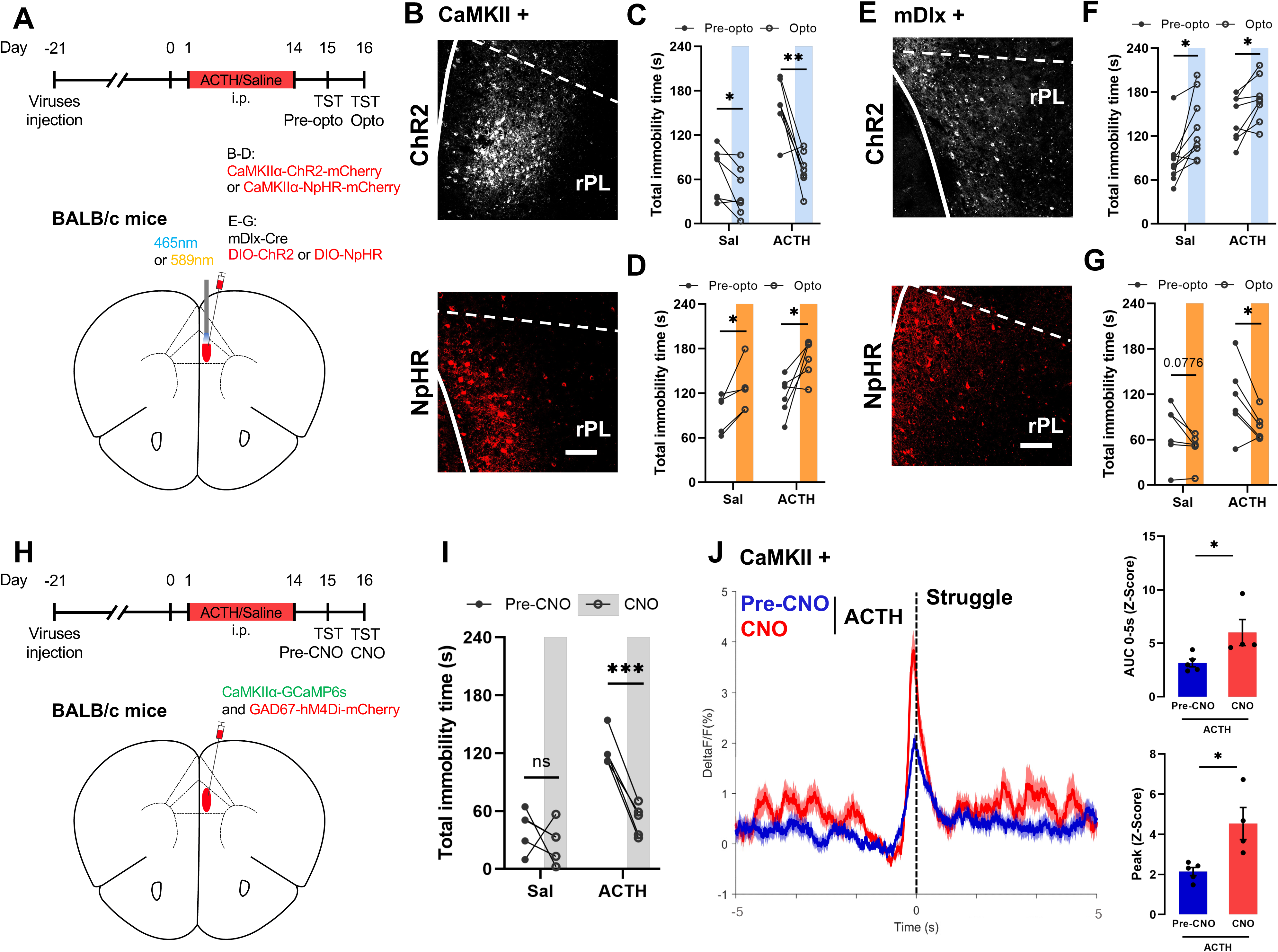
Impact of rPL excitation or inhibition on passive coping behaviour. (A) (Upper) Experimental protocol. (Bottom) Schematic drawing of virus injection site for expressing ChR2 or NpHR in CaMKII+ and mDlx+ neurons, together with optical fiber position. (B, E) Representative image of ChR2-mCherry (upper) or NpHR-mCherry (bottom) expression in CaMKII+ (B) or mDlx+ (E) neurons. Scale bars, 100 μm. (C, D) Total TST immobility time without (Pre-opto, day 15) and with (Opto, day 16) light stimulation to CaMKII+ neurons in mice injected with saline or ACTH. ChR2 [C, *n* = 7 mice/group], NpHR [D, *n* = 5 - 6 mice/group]. (F, G) Total TST immobility time without (Pre-opto, day 15) or with (Opto, day 16) light stimulation to mDlx+ neurons in mice injected with saline or ACTH. ChR2 [F, *n* = 8 – 9 mice/group], NpHR [G, *n* = 6 mice/group]. (H) Experiment protocol (upper) and schematic drawing of virus injection site for expressing GCaMP6s in CaMKII+ neurons and hM4Di in GAD67+ neurons (bottom). (I) Total TST immobility time before and after CNO injection in saline- [*n* = 4] or ACTH-injected [*n* = 4] mice. (J) (Left) Average Ca^2+^ responses (mean ± SEM) in CaMKII+ neuron [*n* = 4] before (blue) and after (red) CNO injection, aligned to the onset (time 0) of struggling in ACTH-injected mice. (Right) Quantifications of AUC (from 0 to 5 s) and peak values of Ca^2+^ responses (pre-CNO vs. CNO). Two-tailed paired *t*-test (C, D, F, G and I). Unpaired *t*-test (J).

In a different set of mice, enhancing inhibitory neuron activity increased total immobility time without inducing rapid p-coping behaviour (Fig. 3F and S-Fig. 4F). In contrast, inhibiting rPL inhibitory neurons shortened the total immobility time and reduced immobility time instantly in ACTH mice, similar to the effect of activating excitatory neurons (Fig. 3G and S-Fig. 4H). In saline-injected mice, inhibiting inhibitory neurons did not reduce passive coping time acutely (S-Fig. 4H), likely due to the fact that the immobility time in these mice was already very low and there was not much room for further reduction. Overall, acutely enhancing rPL excitation reduces p-coping behaviour which leads to immediate struggling, but change in p-coping behaviour emerges slowly when rPL inhibition is enhanced.

The above results on both Ca^2+^ responses and manipulating rPL neuron activity suggest that rPL inhibitory neurons can modulate the activity of excitatory neurons to regulate p-coping behaviour bidirectionally. To determine whether this is the case, we record the Ca^2+^ responses in CaMKII+ neurons while inhibiting rPL inhibitory neurons with the expression of hM4D(Gi) in the GAD67+ neurons, in saline- or ACTH-injected mice (Fig. 3H). Injection of CNO reduced the total immobility time in the ACTH mice, evidenced by higher AUC and peak values in Ca^2+^ responses in the CaMKII+ neurons (Fig. 3I-J). Thus, the rPL inhibitory neurons are a key modulator of p-coping behaviour via their impacts on excitatory neurons.

### Contribution of rPL PV-neurons to passive coping behaviour

Since we found that PV-neurons are the main subtype of inhibitory neurons being affected in chronically stressed mice, there is a possibility that the altered p-coping behaviours in the ACTH mice may be mediated by alterations in the PV-neurons. This requires that PV-neurons are a main modulator of p-coping behaviour. To test this, we monitored PV-neuron activity and manipulated their activities by expressing GCaMP6s, ChR2, or NpHR in the PV-cre mice (Fig. 4A). Since the rate of successfully establishing an ACTH model is very low in the C57BL/6J mice due to their low susceptibility to ACTH, we performed these experiments in non-stressed mice. The Ca^2+^ responses in PV-neurons are time-locked to struggling onset, with similar onset and peak time and onset to peak duration to that in the GAD67+ neurons (Fig. 4B and S-Fig. 5A-E). Blue light pulses not only increased total immobility time but also rapidly increased immobility time, while yellow light instantly decreased immobility time and resulted in a significant reduction of total immobility time (Fig. 4C-E and S-Fig. 5F-G). These results support a rapid and bidirectional modulation of p-coping behaviour by rPL PV-neurons.

**Figure 4.**
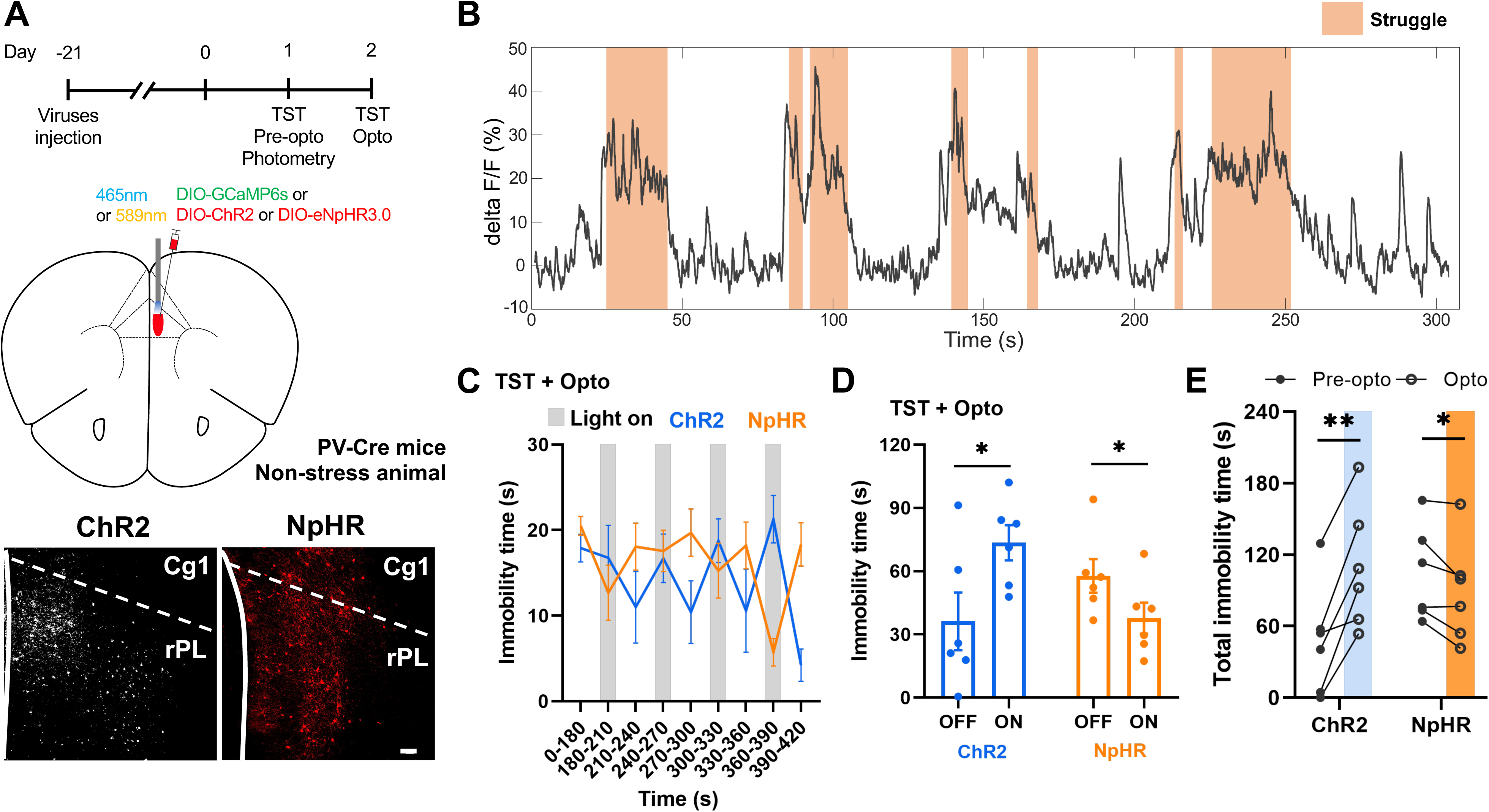
Contribution of rPL PV-neurons to modulations of passive coping behaviour. (A) (Upper) Experimental design and schematic drawing of injection sites for GCaMP6s, ChR2 or NpHR virus in rPL of PV-Cre mice. (Bottom) Representative images of ChR2-mCherry or NpHR-mCherry fluorescence in rPL. Scale bar, 100 μm. (B) Representative traces of Ca^2+^ responses in PV-neurons during TST, with struggling occurred during shaded regions. (C) TST immobility time on day 2. Light illumination began at 180 s, as indicated by grey blocks. (D) Summary of immobility time during light OFF or ON periods, from data in (C). (E) Total immobility time during TST without (Pre-opto, day 1) or with (Opto, day 2) light stimulation in PV-neurons expressing ChR2 [blue, *n* = 6 mice] or NpHR [yellow, *n* = 6]. Unpaired *t*-test (D). Two-tailed paired *t*-test (E).

### (2*R*, 6*R*)-HNK reduces passive coping behaviour

Previous studies have demonstrated that ketamine and its major metabolite (2*R*, 6*R*)-hydroxynorketamine (HNK) rapidly reduces depressive-like behaviours in chronically stressed mice [19]. To understand whether HNK may exert its antidepressant effects by modulating rPL neuron activity, we first examined Ca^2+^ responses in the rPL excitatory and inhibitory neurons during TST in chronically stressed mice 1 hour after HNK injection (Fig. 5A). We found that HNK injection significantly reduced immobility time during TST (Fig. 5B-C). This reduction was associated with a reduction in the immobility episode duration but not number (Fig. 5D). Compared to before HNK injection (Pre-HNK), Ca^2+^ responses in the CaMKII+ neurons were of higher AUC and peak values after HNK injection (Fig. 5E). A reduction in the AUC of Ca^2+^ responses was observed in the GAD67+ neurons (Fig. 5F and S-Fig. 6A). These changes collectively led to a higher E/I ratio in rPL (Fig. 5G). In addition, HNK injection resulted in reduced onset to peak of Ca^2+^ responses in the GAD67+ neurons and a longer increase in Ca^2+^ responses in CaMKII+ neurons, leading to a stronger excitation before struggling onset (Fig. 5H and S-Fig. 6B-C).

**Figure 5.**
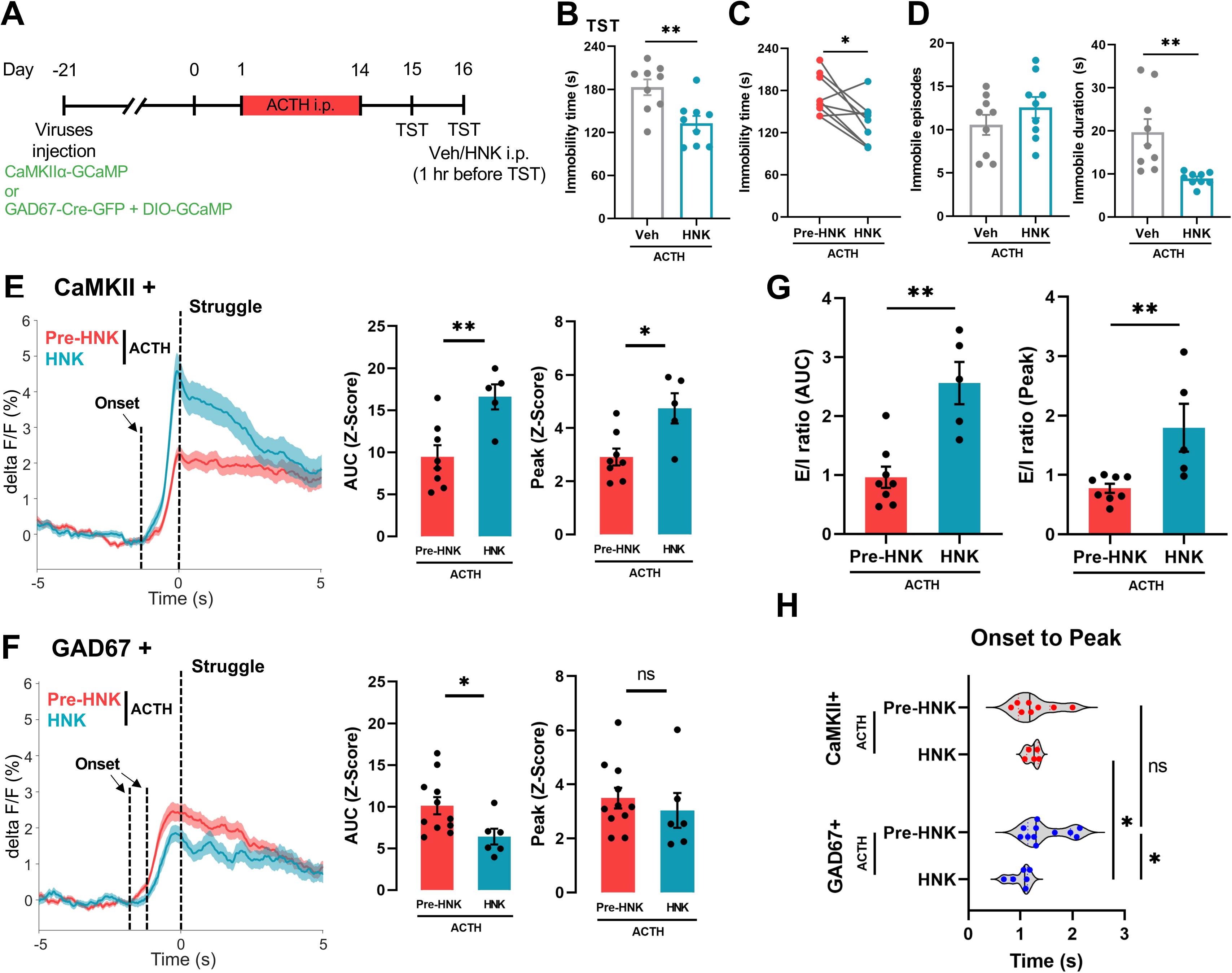
HNK reduces passive coping behaviour in chronically stressed mice. (A) Experimental protocol. (B) Immobility time in ACTH-mice 1 hour after injection with vehicle [Veh, *n* = 9] or HNK [*n* = 9]. (C) Immobility time before (day 15) and after (day 16) HNK [*n* = 9] injection in the same set of ACTH-injected mice. (D) Numbers of immobility episodes (left) and averaged duration of each immobility episode (right) during TST in Veh- and HNK-injected mice. (E, F) (Left) Averaged Ca^2+^ responses (mean ± SEM) in CaMKII+ (E) or GAD67+ (F) neurons in Pre-HNK [*n* = 8 - 11] and HNK [*n* = 5 - 6] mice, aligned to struggling onset (time 0). (Right) Quantifications of AUC (from 0 to 5 s) and peak values. (G) E/I ratio calculated from Ca^2+^ responses by dividing AUC (left) or peak values (right) of CaMKII+ neurons to that in GAD67+ neurons (Pre-HNK vs. HNK). (H) Quantification of duration in Ca^2+^ responses (from response onset to peak) in CaMKII+ or GAD67+ neurons, before and after HNK injection. Unpaired *t*-test (B, D, E, F, G and H). Two-tailed paired *t*-test (C).

### Impacts of (2*R*, 6*R*)-HNK on rPL excitatory and inhibitory neurons in chronically stressed mice

Since we have observed altered rPL excitatory and inhibitory neurons activity and properties in chronically stressed mice, and demonstrated that both of them modulate p-coping behaviour, we next examined whether HNK may affect rPL excitatory and inhibitory neurons to restore p-coping behaviour. First, we recorded spike rates in the rPL excitatory and inhibitory neurons *in vivo* (S-Fig. 6D), and we observed a higher spike rate in excitatory neurons and a lower spike rate in inhibitory neurons, starting ∼ 30 min after HNK injection and lasted for at least 24 hours (Fig. 6A). These opposite changes led to a higher rPL E/I ratio (Fig. 6B). Next, we recorded from rPL neurons in slices 1 hour after injection of either vehicle or HNK in ACTH-injected mice. HNK did not alter either pre- or post-synaptic excitatory transmissions in the rPL excitatory neurons participating in TST, but HNK significantly increased the intrinsic excitability of these neurons (Fig. 6C-D and S-Fig. 6E). In comparison, HNK significantly reduced the frequency, amplitude and area of sIPSC in the excitatory neurons (Fig. 6E). These changes resulted in a higher E/I ratio (Fig. 6F). In the FSNs, HNK reduced sEPSC frequency (Fig. 6G) but not sEPSC amplitude, area or decay time (S-Fig. 6F), together with a significant reduction in their intrinsic excitability (Fig. 6H). These findings collectively suggest a reduced activity in the FSNs. In contrast, HNK had no significant effect on any of the parameters measured on RSNs (Fig. 6I-J and S-Fig. 6G). Altogether, HNK partially reverses the chronic stress-induced alterations in the rPL neurons and induces additional inhibitory alterations.

**Figure 6.**
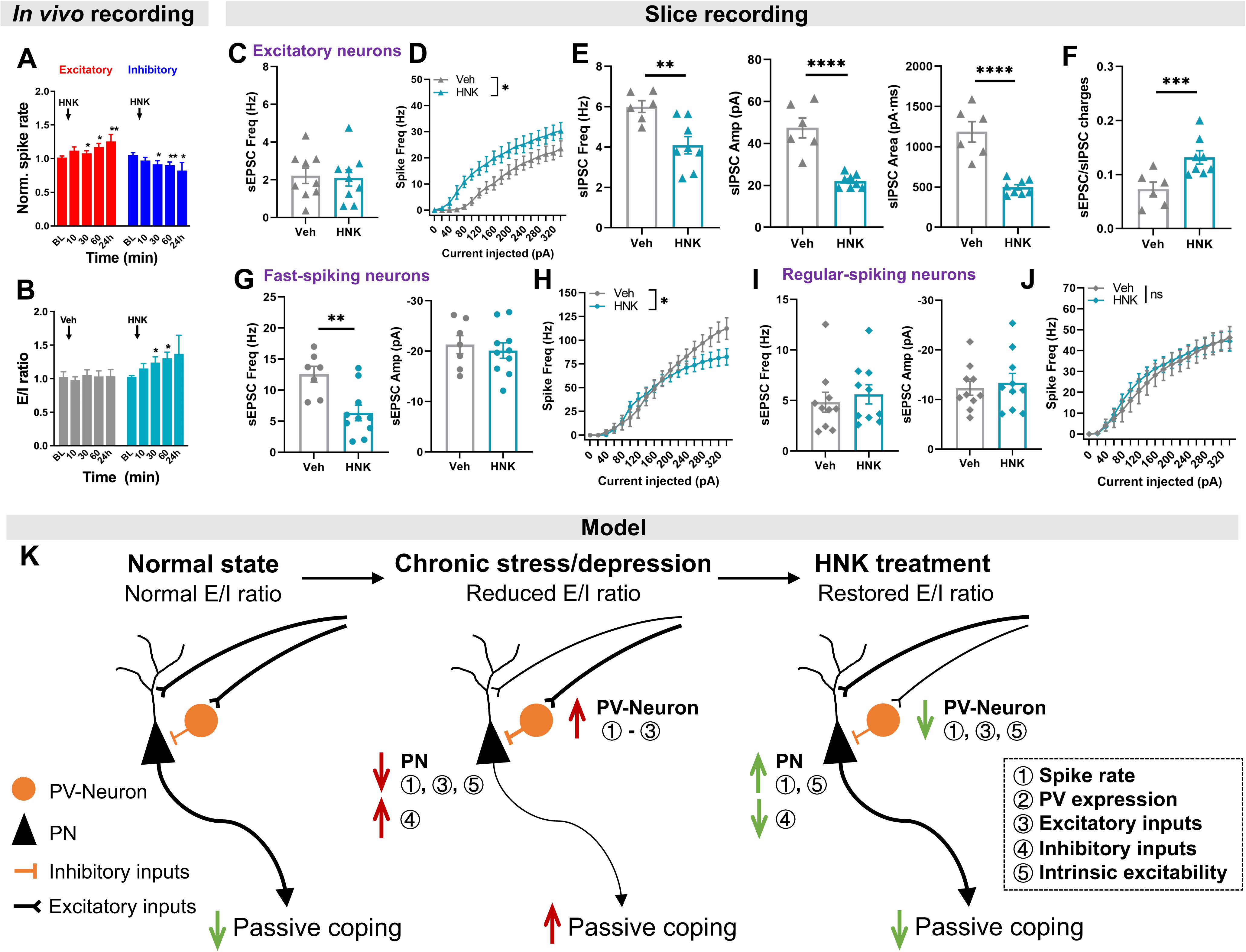
HNK partially reverses synaptic alterations in rPL neurons in chronically stressed mice. (A) Time course of changes in *in vivo* spike rate in excitatory [red, *n* = 201 cells/10 mice] or inhibitory [blue, *n* = 84 cells/12 mice] neurons after HNK injection. (B) E/I ratio of *in vivo* spike rate calculated in ACTH-mice treated with Veh or HNK. (C) Quantification of sEPSC frequency in excitatory neurons from ACTH-mice treated with Veh [*n* = 9 cells/3 mice] or HNK [*n* = 9 cells/3 mice]. (D) Intrinsic excitability in excitatory neurons from ACTH mice treated with Veh [*n* = 9 cells/3 mice] or HNK [*n* = 9 cells/3 mice]. (E) Quantification of sIPSC frequency, amplitude, and area in excitatory neurons from ACTH-mice treated with Veh [*n* = 6 cells/2 mice] or HNK [*n* = 8 cells/2 mice]. (F) E/I ratio of sEPSC charges/sIPSC charges calculated by dividing total area of sEPSC over sIPSC in the same excitatory neurons. (G, H) Quantification of sEPSC frequency and amplitude (G), and intrinsic excitability (H) in fast-spiking GAD67+ neurons from ACTH-mice treated with Veh [*n* = 7 cells/3 mice] or HNK [*n* = 10 cells/3 mice]. (I, J) Quantification of sEPSC frequency and amplitude (I), and intrinsic excitability (J) of regular-spiking GAD67+ neurons from ACTH-mice treated with Veh [*n* = 10 cells/3 mice] or HNK [*n* = 10 cells/3 mice]. (K) A model summarizing the major findings. Two-way ANOVA (A, B, D, H and J). Unpaired *t*-test (C, E, F, G, and I).

## DISCUSSION

In this study, we demonstrated that rPL inhibitory neurons, especially PV-neurons, have a profound impact on the rPL excitatory neurons to modulate p-coping behaviour. Chronic stress enhances the activity of PV-neurons via alterations in both synaptic and neuronal properties, and prolongs p-coping behaviour. Fast-acting antidepressant HNK rapidly restores the p-coping behaviour, partially reverses the chronic stress-induced rPL alterations, and appears to exert its effect via modulation of rPL E/I ratio. These findings highlight the importance of rPL inhibition in the modulation of p-coping behaviour, and may provide new targets for better treatment of depression.

Consistent with previous reports [8, 9, 17], we found that activation of rPL E-neurons elicits a rapid termination of p-coping behaviour (i.e., immobile period during TST), and this observation emphasizes the importance of rPL excitation in the regulation of p-coping. The excitation-induced rapid emergence of struggling switches the coping strategy from passive to active. In contrast to p-coping, active coping allows an animal to escape from a stressor or stressful circumstance and hence may eliminate the source of stress altogether [27]. HNK elicits a rapid increase in the rPL E-neuron activity, associated with a rapid reduction in p-coping behaviour, which may rapidly relieve certain symptoms in depression patients.

In contrast to the E-neurons, rPL I-neurons inhibit E-neurons and prolong the immobility period and hence enhance p-coping behaviour. This action is slow since optogenetic activation of I-neuron activity does not lead to rapid changes in TST immobility but rather a slow emergence of such behaviour over time. These observations suggest that rPL I-neurons sustain the status quo, perhaps to conserve energy (see below), until a better opportunity to switch to active coping. Whether this is the case requires future study to explore. HNK partially reverses chronic stress-induced synaptic transmission in rPL PV/FSNs, and reduces intrinsic excitability of PV/FSNs, with the net effect of reducing the activity in I-neurons. Together with higher excitation induced by HNK, the net result is a restoration of the E/I ratio to the level in the non-stressed mice. These findings provide the cellular basis for HNK’s fast antidepressant effect in PFC. Collectively, our findings emphasize that rPL E/I balance (ratio) is a critical factor in the generation/modulation of p-coping behaviour. This conclusion is consistent with Yoon, et al. [28] and findings in humans. For example, meta-analyses of ^1^H-MRS studies in MDD patients reported reduced levels of glutamate and Glx (a composite of glutamate + glutamine) in the dorsolateral PFC and anterior cingulate cortex, which is associated with MDD diagnosis and severity [29–32].

As the rPL E/I ratio is lower in chronically stressed mice, there are two ways to restore this: to enhance E or to reduce I. We have observed that HNK mostly engages the former mechanism while optogenetic inhibition of PV-neurons targets the latter, and both are effective in reverting the E/I ratio and p-coping behaviour. Similar findings revealed that GABAergic neurons are an essential target to mediate the antidepressant-like effects (reduced immobility in FST) as this effect was blocked by chemogenetic activation of Gad1+, somatostatin+, and PV+ neurons [16, 33, 34]. The mechanism of HNK we have revealed here is consistent with HNK not affecting NMDAR activity at its antidepressant dose (i.p. 10 mg/kg, roughly ∼ 8 μM in the brain) [35]. This is in clear contrast to HNK’s parent compound, ketamine, which is a potent NMDAR inhibitor and known to affect I-neuron activity [16, 33, 36–38].

The rPL PV-neurons appear to be the main inhibitory neuron subtype that modulates p-coping via their powerful perisomatic synaptic inhibition onto other neurons (mostly E-neurons) [24, 39]. Importantly, chronic stress induced by ACTH injection mostly affects the rPL PV-neurons while reversing (i.e., reducing) rPL PV-neuron activity reversed the altered p-coping behaviour in the stressed mice (Fig. 4C-E). Both increased and decreased activity of PV/I-neurons have been reported in different models and it is at least partially related to the duration of stress administered in that short-term (2 - 4 weeks) leads to enhanced I-neuron activity while long-term (5 - 9 weeks) results in reduced function [40]. Interestingly, HNK also selectively affects PV-neurons to regulate rPL excitation in stressed mice. Since we only examined the HNK effect 1 hour after injection, we do not know whether HNK may exert a larger or longer effect on PV-neurons but we have shown that HNK’s effect on inhibitory neurons in rPL lasts at least 24 hr (Fig. 6A). Notably, previous studies have reported that the PFC regular-spiking inhibitory neurons participate in the top-down regulation of hippocampal information processing [41], but the alteration in stress model was poorly understood. Here, we found no effects on RSN in response to chronic stress and HNK. Previous studies have shown altered SOM-neuron functions in depression models and modulating SOM-neuron activity contributes to the anti-depressant effects [16, 33, 34]. It is possible that distinct subpopulations of inhibitory neurons may participate in the chronic stress-induced alterations which may at least partially contribute to the heterogeneity of depression.

We propose that p-coping behaviours, on a short-term basis, may be beneficial for the organism to avoid persistently consuming energy in face of situations where stressors/stress environments cannot be removed/avoided [2]. A significant reduction in the astrocyte structure and functions has been reported in both humans and animals, including MDD patients [42–46]. As astrocytes are a major energy contributor for neurons [47], this deficit is consistent with reduced energy expenditure in the brain under chronic stress [4, 43, 48]. Switching from p-coping to a-coping requires higher energy usage, and whether HNK may provide such required energy remains future investigation.

In summary, chronic stress and HNK mostly affected fast-spiking/parvalbumin inhibitory neurons rather than other inhibitory neurons, indicating the important role of this subtype of inhibitory neurons. These neurons also modulate E/I balance to regulate passive coping that is influenced by chronic stress and rapidly restored by fast-acting antidepressant.

## MATERIAL AND METHODS

### Animals

Male BALB/c, BALB/c.GAD67-GFP and C57BL/6J.PV-Cre mice used were bred and group-housed (5-6 mice/cage) in the Laboratory Animal Center of Shenzhen Graduate School, Peking University. The animals were subjected to standard laboratory conditions (22℃ ± 2℃, 50% - 70% humidity, and a 12 hr/12 hr light/dark cycle) and food and water were provided *ad libitum*. Bedding, water, and food were changed once a week. Male mice of 9-16 weeks of age were used for chronic stress model establishment, behavioural tests, as well as *in vivo* electrophysiology and fiber photometry recordings. All of the *in vivo* experimental procedures were performed in accordance with the ARRIVE (Animal Research: Reporting of In Vivo Experiments) guidelines. All of the protocols used for animals were approved by the Peking University Shenzhen Graduate School.

### Chronic stress model by ACTH injection

Male BALB/c mice were intraperitoneally injected with adrenocorticotropic hormone (ACTH, 1 mg/kg) for 14 successive days to establish the chronic stress model. Concurrently, male BALB/c mice injected with saline were set as the control group to ACTH-injected mice.

### Virus injection and optical fiber implantation

To record Ca^2+^ responses in the excitatory and inhibitory neurons in rPL (2.2 mm anterior to Bregma; ± 0.3 mm lateral to midline; 2.2 mm below skull surface), rAAV2/9-CaMKIIα- GCaMP6s-WPRE-hGH-pA (virus titer: 5.21E+12 vg/ml) and rAAV2/9-GAD67-Cre-EGFP-WPREs (4.78E+12 vg/ml) mixed with rAAV2/9-Ef1α-DIO-GCaMP6s-WPRE-pA (5.54E+12 vg/ml, ratio: 1:1) were injected, respectively. To optogenetically manipulate excitatory neurons in rPL, mice were injected with rAAV2/9-CaMKIIα-hChR2(H134R)-mCherry (3.34E+12 vg/ml) or rAAV2/9-CaMKIIα-eNpHR3.0-mCherry (4.48E+12 vg/ml). To optogenetically manipulate inhibitory neurons in rPL, mice were injected with rAAV2/9-mDlx-SV40 NLS-Cre (5.15E+12 vg/ml) mixed with rAAV2/9-Ef1α-DIO-hChR2(H134R)-mCherry (5.54E+12 vg/ml, 1:1) or rAAV2/9-Ef1α-DIO-eNpHR3.0-mCherry (3.34E+12 vg/ml, 1:1). To chemogenetically manipulate inhibitory neurons in rPL, mice were injected with rAAV2/9-rGAD67-hM4D(Gi)-EGFP (3.12E+12 vg/ml). To label the neurons that selectively participated in TST, mice were injected with rAAV2/9-cFos-tTA-WPRE-hGH-pA (5.93E+12 vg/ml) mixed with rAAV2/9-TRE3g-mCherry-WPRE-hGH-pA (5.07E+12 vg/ml, 1:1) and were fed by food with doxycycline (DOX, 40 mg/kg).

Following the virus injection, a fiber optic cannula (2 mm length, 200 μm diameter, NA = 0.37, 1.25 mm Ferrule Size, Inper LLC) was inserted towards the virus-targeted region in rPL for mice subjected to *in vivo* imaging and opto-manipulation. The ceramic ferrule was supported with two skull-penetrating screws and dental cement.

### Fiber photometry recording

Mice injected with GCaMP6s virus were connected to a mono fiber optic (200 μm diameter, NA = 0.37, 1.5 m long) through ceramic ferrule for fluorescence transmission from the targeted cells. The excitation LED light (470 nm) was reflected via a dichroic mirror (MD498; Thorlabs) and focused through a 20×/0.4 NA objective lens. Fluorescence signal of GCaMP6s was acquired at a sampling rate set to 50 frames/s using a multichannel fiber photometry system (ThinkerTech Nanjing Bioscience Inc., China) and was band-pass filtered (MF525-39) before collection using a CMOS (DCC3240M, Thorlabs). The power of light (20-25 μW) was measured and adjusted to minimize the bleaching effects from the tip of optical fiber. MATLAB was utilized for data analysis. Quantification of fluorescence values (ΔF/F) was calculated and expressed in z-score. The magnitudes of Ca^2+^ responses were measured from the area under the curve (AUC) and the peak value.

### Behavioural testing

Tail suspension test (TST): 1 hr prior to the test, mice were placed in the test room to habituate the environment. The TST apparatus is an opaque rectangular box (height: 50 cm, length and width: 15 cm) with one horizontal crossbar on top of the box and one side opening for camera recording the mice motion. The distal fourth of the mouse’s tail was attached to the crossbar by medical adhesive tape and the mouse was suspended upside-down to approximately half of the box height, ensuring that the mice do not struggle against the box wall. Mice were tail-suspended for a total of 7 min and the last 4 min of the test were available for quantification. The motion of mice was analysed by ANY-maze (Version 7.08, U.S.). The immobility time was defined as the duration of mice completely immobile or with slight swing or with small movement of the forelimbs. The apparatus was then cleaned with 75% ethanol between trials to prevent odor interference. For optogenetic manipulation, light stimulation was given starting from the 3 min of the test for 30 s, followed by 30 s intervals, repeating 4 times of light stimulating-interval sections till the end of the test. For chemogenetic manipulation, CNO was injected 30 min before the test.

Forced swimming test (FST): Mice were forced to swim in a Plexiglas cylinder (height: 50 cm, diameter: 20 cm) containing water (24℃ ± 1℃) to a depth of 30 cm. A cardboard was placed between the cylinders to prevent disturbance. Mice were forced-swam for a total of 7 min and the last 4 min of the swimming were available for quantification. A camera acquired the swimming process and the trajectory of mice was analysed by EthoVision XT (Noldus, Netherlands). The immobility time was defined as the duration of mice floating in a motionless state or with slight movement to prevent sinking. Water was replaced between tests to avoid odor interference. At the end of the test, mice were placed on a heating pad to restore body temperature.

Sucrose preference test (SPT): 24 hr before the test, mice were trained to habituate to the 1% sucrose solution (weight/volume) and the location of the bottles. One bottle of 1% sucrose solution and one bottle of tap water were placed in the cage and their location was swapped after 12 hr. Mice were then deprived of water for 12 hr. Subsequently, mice were individually housed in a cage with two bottles with 100 ml 1% sucrose solution or tap water for 24 hr. Bottles were swapped after 12 hr to avoid place preference. At the end of the test, sucrose preference was calculated as the percentage of the consumed volume of sucrose solution relative to the total liquid intake.

### Electrophysiological recording on slices

Mice were anesthetized using 1% pentobarbital sodium in saline followed by decapitalization. Dissected brains were rapidly placed in and ice-cold oxygenated (95% O_2_ and 5% CO_2_) slicing solution containing (in mM) 110 choline chloride, 7 MgSO_4_, 2.5 KCl, 1.25 NaH_2_PO_4_, 25 NaHCO_3_, 25 D-glucose, 11.6 sodium ascorbate, 3.1 sodium pyruvate, and 0.5 CaCl_2_. Coronal sections (300 μm) containing rPL were obtained using a Leica Vibratome (VT-1200S, Leica Microsystems, Wetzlar, Germany) and firstly incubated for 30 minutes at 35°C for recovery then 1 hr at room temperature (25-30°C) with oxygenated artificial cerebrospinal fluid (aCSF) containing (in mM): 127 NaCl, 2.5 KCl, 1.25 NaH_2_PO_4_, 25 NaHCO_3_, 25 D-glucose, 2 CaCl_2_, and 1 MgCl_2_.

Excitatory neurons selectively labeled in TST by c-Fos from rPL, and inhibitory neurons indicated by GFP fluorescence in GAD67-GFP mice were identified on slices through a microscope (BX51WI, Olympus, Japan) with a 40× water-immersion differential interference contrast objective. Brain slices under the microscope were constantly perfused (2.0 mL/min) in oxygenated aCSF at room temperature. Neurons were recorded using HEKA EPC10 double patch clamp amplifier (HEKA Elektronik) at a sampling rate of 10 kHz and filtered at 2 kHz. Recording pipettes (5-7 MΩ) were filled with K^+^-based intracellular solution containing (in mM) 128 K-Gluconate, 10 NaCl, 2 MgCl_2_, 0.5 EGTA, 10 HEPES, 0.4 Na_2_GTP, and 4 Na_2_ATP. For sIPSC recording, recording pipettes were filled with intracellular solution containing (in mM) 125 CsMeSO_5_, 5 NaCl, 1.1 EGTA, 10 HEPES, 0.3 NA_2_GTP, 4 Mg-ATP, and 5 QX-314.

### *In vivo* recording

Male mice aged 9-16 weeks were firstly anesthetized with isoflurane (induction 3% and maintenance 1.5%) and placed on a stereotaxic apparatus. Skull was exposed for two screw implantation to secure electrode array implants. mPFC (2.1 mm anterior to Bregma; ± 0.3 mm lateral to midline; 2.2 mm below skull surface) was targeted by multiwire electrodes which were unilaterally implanted. The electrodes are composed of 16 nichrome wires (individually insulated, 35 μm inner diameter, impedance 300 ∼ 900 KΩ; Stablohm 675, California Fine Wire) in a 3×5×5×3 arrangement (spacing between wires ∼ 200 μm). Electrodes were attached to a connector (Mil-Max) and secured with dental cement.

Animals were allowed 7-14 days to recover from surgery and baseline neural signals were recorded before ACTH/Saline injection or HNK/Veh injection. Following ACTH/Saline injection every day, mice were recorded for 15 min in their home cage from day 1 to day 14. ACTH-injected mice were recorded (15 min) after HNK/Veh injection at 10, 30, 60 minute and 24 hour in the home cage. Broadband (0.3 Hz to 7.5 kHz) neural signals from the implanted 16-channel arrays were simultaneously recorded (16 bits at 30 kHz) by a 64-channel acquisition system (Zeus, Bio-Signal Technologies). Acquired data (i.e., extracellular spikes) were aligned and sorted (offline sorter software, Plexon). Sorted spikes were further analysed using NeuroExplorer (Nex Technologies), and identified as excitatory/inhibitory neurons by categorizing their peak-to-trough and half-width values.

### Immunofluorescence and imaging

Mice were anesthetized by overdose pentobarbital sodium and underwent transcardial perfusion with 20 mL phosphate-buffered saline (PBS) before 20 mL 4% paraformaldehyde (PFA). Mouse brains were then collected and fixed overnight by PFA at 4℃, followed by 30% sucrose in PBS for 48 hours. Coronal sections containing rPL (30 μm) were sliced using a cryostat microtome (Leica, Wetzlar, Germany) and were washed for 15 min in 0.5% Triton X-100 in PBS (PBST). Slices were then incubated with 10% normal goat serum (NGS) in PBST for 1.5 hr, followed by primary antibody (Parvalbumin Rabbit mAb, 1:1000, #80561, Abcam) in PBST + 10% NGS at 4℃ overnight. The incubated slices were then washed by PBS for two times (15 min/each) and incubated with Alexa Fluor^TM^ 647 donkey anti-rabbit IgG (1:1000, A31573, Invitrogen) in PBST for 2 hr at room temperature. Slices were stained by DAPI (1:1000, Beyotime, China) for nuclei counterstain for 10 min and washed for 15 min in PBS. A confocal microscope (A1R, Nikon, Japan) was utilized for sections imaging.

### Statistics

All statistical analyses were performed with GraphPad Prism 9 for Windows (Version 9.5.0). Figures were generated using Adobe Illustrator 2021. Data are shown as mean ± SEM. Statistical significance was expressed as *, *p* < 0.05; **, *p* < 0.01; ***, *p* < 0.001; ****, *p* < 0.0001; ns, not significant.

## ACKNOWLEDGEMENTS

This work is supported by grant from Shenzhen-Hong Kong Institute of Brain Science-Shenzhen Fundamental Research Institutions (2023SHIBS0004) and grants (82471548, Y2023064). We thank the members of Zhou’s lab for the technical supports and helpful discussion.

## AUTHOR CONTRIBUTIONS

Conceptualization: QZ, THF, and XM. Methodology: QZ, THF, TL, XM, and XC. Experiment and data analysis: THF, TL, and QZ. Writing – original draft: QZ and THF. Writing – review and editing: QZ, THF, and XC. Funding acquisition: QZ and XC. Project administration: QZ. Supervision, QZ.

## DATA AVAILABILITY

All data are available in the main text or supplementary materials upon request.

## ADDITIONAL INFORMATION

Supplementary information consisted of supplementary figure 1-6

