## Supplementary material for "Key contribution of prefrontal inhibition to passive coping behaviour: chronic stress and fast-acting antidepressant": Fig S1-6

### Supplementary Figure 1

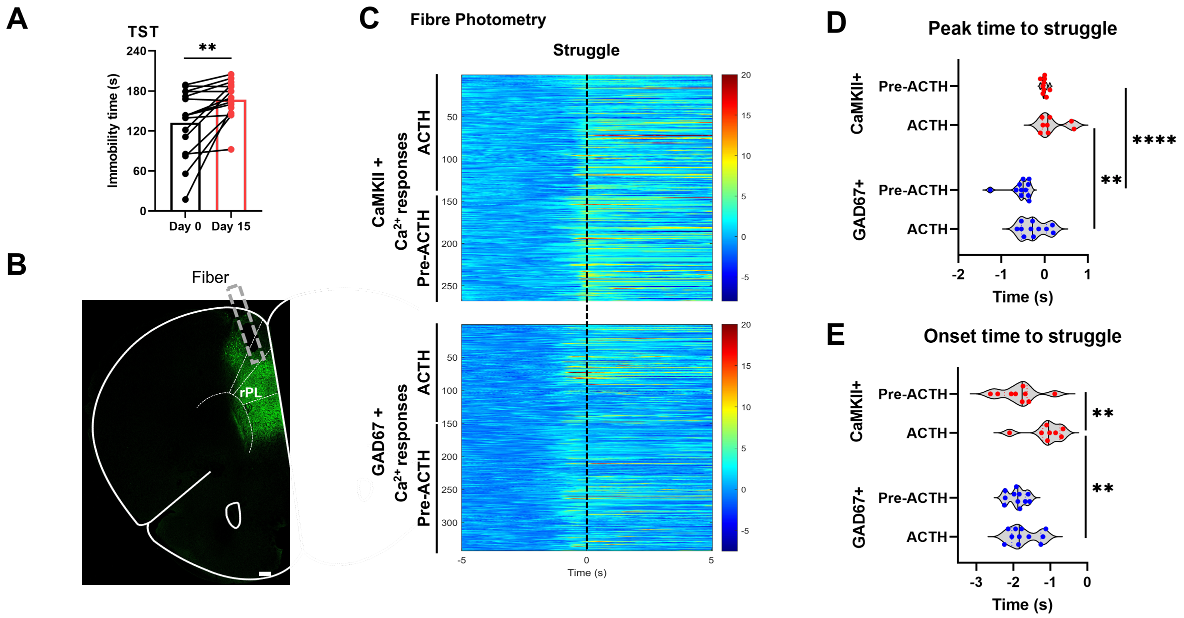


**S-Figure 1. Properties of rPL $Ca^{2+}$ responses during TST.**

(A) Immobility time during TST before and after ACTH injection in the same set of mice [$n$ = 16 mice].

(B) Sample image of GCaMP6s expression with location of optical fiber (grey dotted rectangle) in BALB/c mice. Scale bar, 200 μm.

(C) Heatmaps of $Ca^{2+}$ responses in CaMKII+ and GAD67+ neurons before and after ACTH injection in the same set of mice. Responses aligned to the onset of struggling (time 0).

(D) Quantification of peak to struggle onset (0 s) in $Ca^{2+}$ responses for CaMKII+ and GAD67+ neurons.

(E) Quantification of onset to struggle onset (0 s) in $Ca^{2+}$ responses for CaMKII+ and GAD67+ neurons.

Two-tailed paired $t$-test (A). Unpaired $t$-test (D and E). *, $p < 0.05$; **, $p < 0.01$; ***, $p < 0.001$; ****, $p < 0.0001$; ns, not significant.

**Supplementary Figure 2**

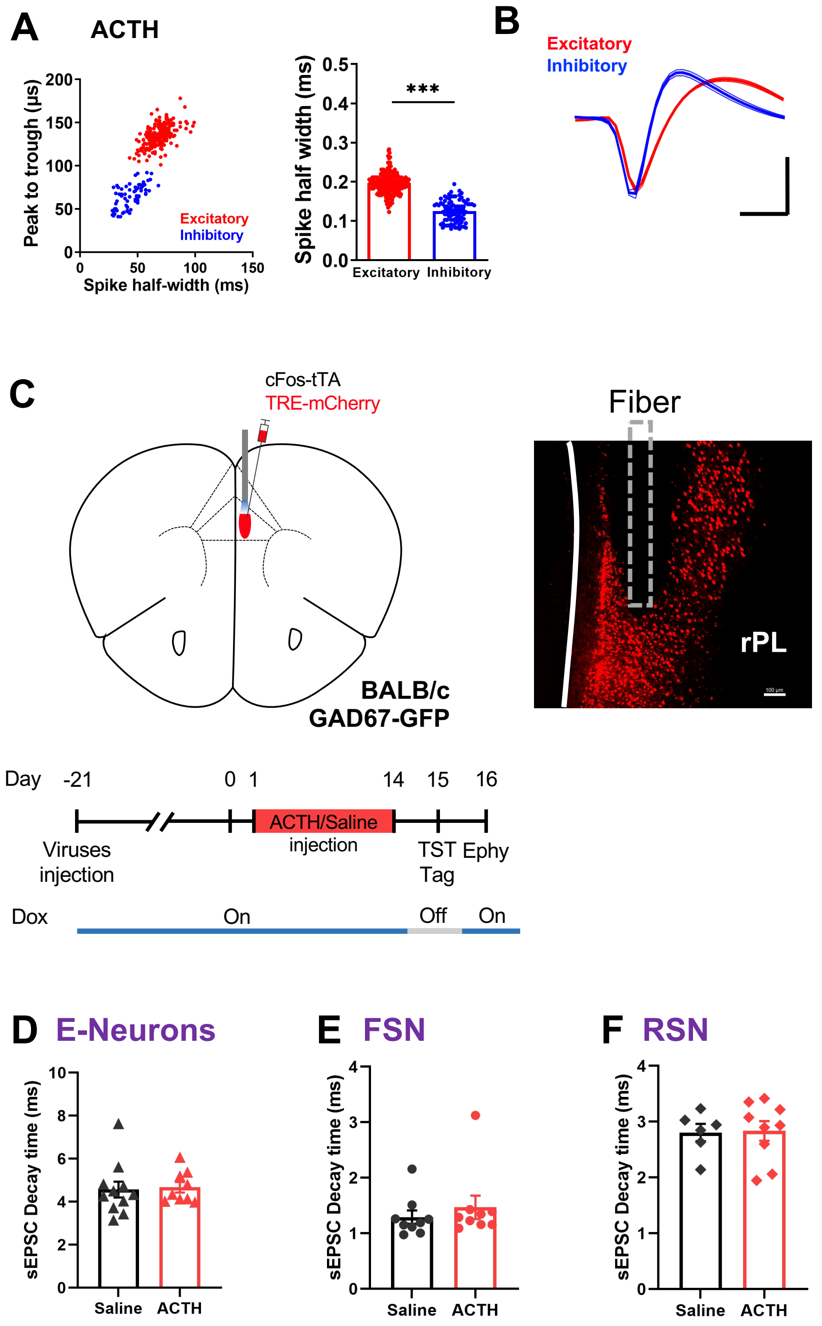


**S-Figure 2. Electrophysiological properties of recorded rPL excitatory and inhibitory neurons.**

(A) Identification of excitatory [$n$ = 220 cells/10 mice] and inhibitory [$n$ = 83 cells/10 mice] neurons from *in vivo* recordings in ACTH-injected mice, based on peak-to-trough and spike half-width values.

(B) Averaged sample spike waveforms of putative excitatory and inhibitory neurons recorded *in vivo*. Scale bars, 50 μV and 250 μs.

(C) (Upper) Site for virus injection and sample image of c-Fos labelled cells/optical fiber location (grey dotted rectangle) in BALB/c mice. (Bottom) Experiment protocol. Scale bar, 100 μm.

(D) sEPSC decay time in c-Fos labelled excitatory neurons (E-Neurons) in rPL slices.

(E) sEPSC decay time in fast-spiking GAD67+ neurons (FSN) in rPL slices.

(F) sEPSC decay time in regular-spiking GAD67+ neurons (RSN) in rPL slices.

Unpaired *t*-test (A, D, E and F).

### Supplementary Figure 3

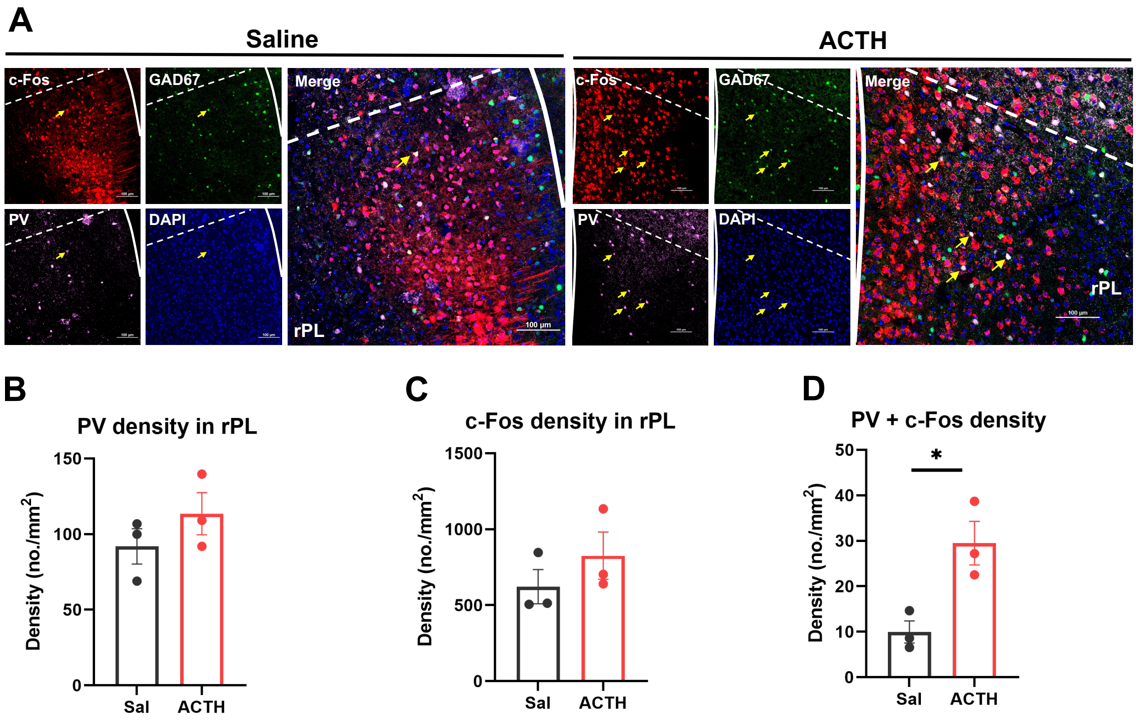


**S-Figure 3. Higher rPL inhibition associated with TST in chronically stressed mice.**

(A) Representative images of rPL sections showing GAD67-GFP (Green), PV immunofluorescence (Violet), and TST-associated c-Fos-positive (Red) neurons in saline- or ACTH-injected mice [$n$ = 3 mice/group]. Blue, DAPI staining. Arrows indicate neurons with overlapping fluorescence from all channels. Scale bar, 100 µm.

(B) Density of PV-positive neurons in rPL.

(C) Density of c-Fos-positive neurons in rPL.

(D) Density of rPL neurons positive for both PV and c-Fos.

Unpaired $t$-test (B - D).

**Supplementary Figure 4**

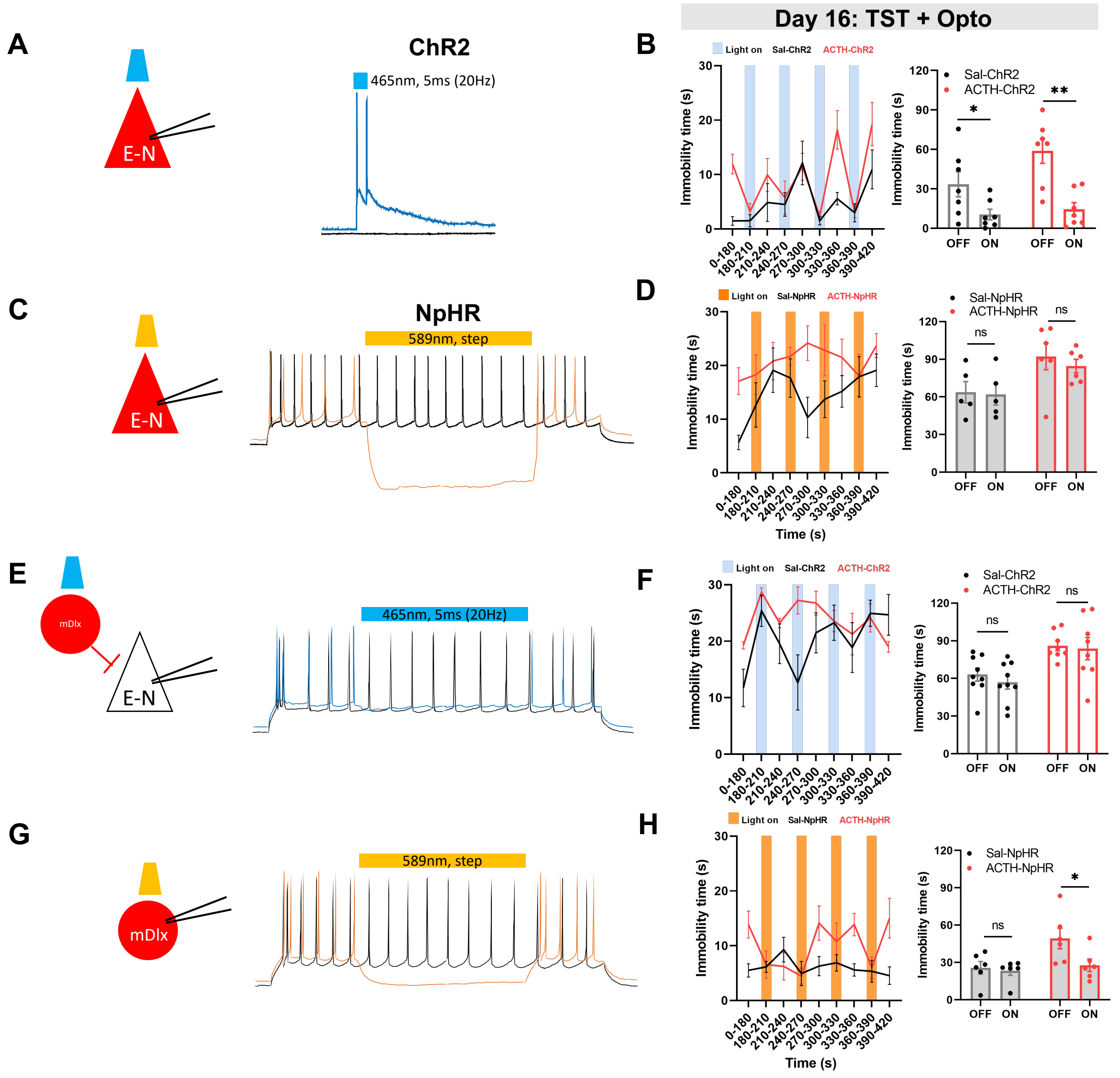


**S-Figure 4. Acute manipulation of rPL excitatory and inhibitory neurons.**

(A, C) Light activation (A) or inhibition (C) of CaMKII+ excitatory neurons (E-N) in rPL slices.

(B, D, F, H) (Left) Quantification of TST immobility time with opto-manipulation on day 16. Light activation or inhibition began at 180 s as indicated by blue or yellow blocks, respectively. (Right) Summary data on light OFF vs. ON from the left panels. Optogenetic manipulation for CaMKII+ neurons expressing ChR2 [B, $n$ = 7/group] or NpHR [D, $n$ = 5-6/group], and mDlx+ neurons expressing ChR2 [F, $n$ = 8-9/group] or NpHR [H, $n$ = 6/group].

(E) Impact of light activation of mDlx+ neurons on excitatory neurons, recorded in rPL slices

(G) Light inhibition of mDlx+ neurons in rPL slices.

Unpaired *t*-test (B, D, F and H).

### Supplementary Figure 5

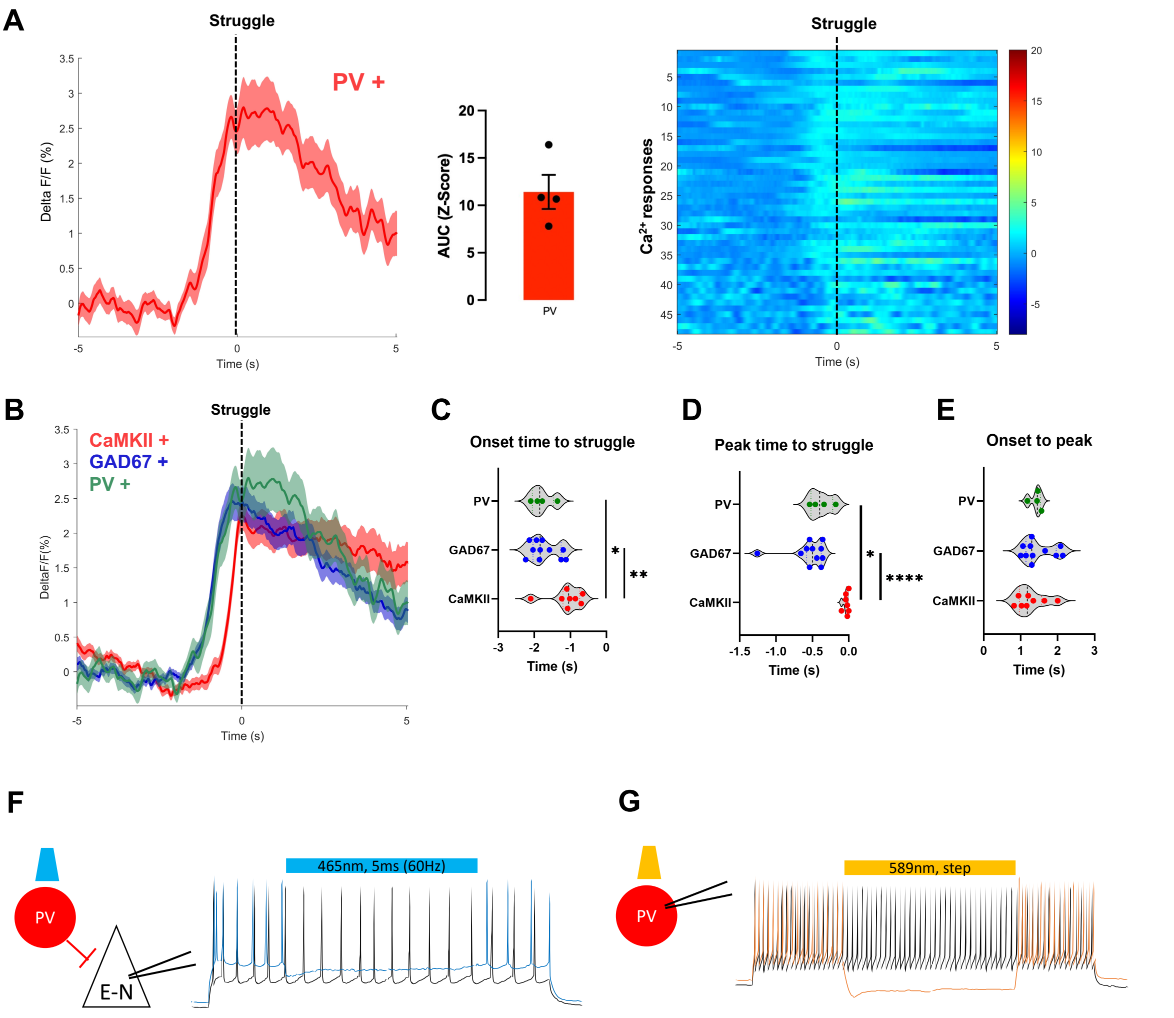


**S-Figure 5. rPL PV-neuron responses during TST.**

(A) Averaged representative traces (mean ± SEM, left) and heatmaps (right) of $Ca^{2+}$ responses in PV-neurons aligned to struggling onset. AUC (from 0 to 5 s, middle) [$n = 4$].

(B) Averaged $Ca^{2+}$ responses (mean ± SEM) in PV+ (Green), GAD67+ (Blue), and CaMKII+ (Red) neurons aligned to struggle onset (time 0).

(C) Quantification of onset to struggle onset (0 s) in $Ca^{2+}$ response for PV+, GAD67+ and CaMKII+ neurons.

(D) Quantification of peaks to struggle onset (0 s) in $Ca^{2+}$ response for PV+, GAD67+ and CaMKII+ neurons.

(E) Quantification of onset to peak duration in $Ca^{2+}$ responses for PV+, GAD67+ and CaMKII+ neurons.

(F) Impact of light activation of PV-neurons on excitatory neurons (E-N), recorded in rPL slices.

(G) Light inhibition of PV-neurons in rPL slices.

Unpaired *t*-test (C - E).

### Supplementary Figure 6

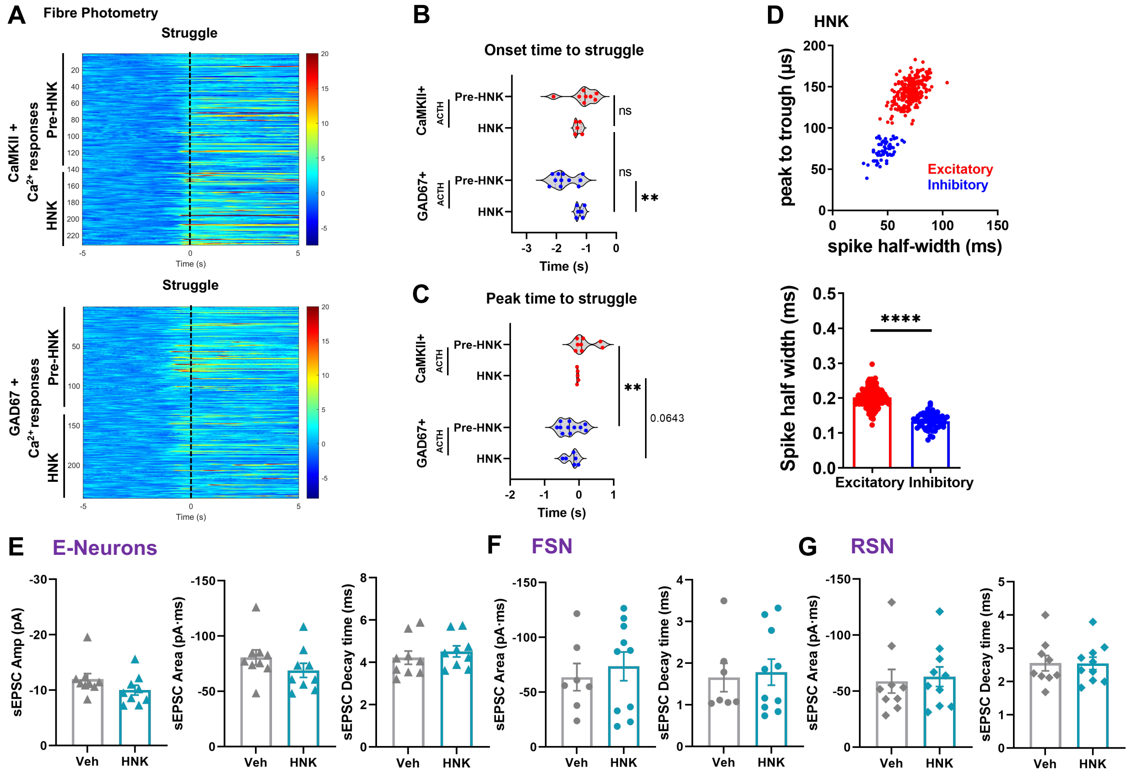


**S-Figure 6. HNK effects on rPL excitatory and inhibitory neurons.**

(A) Heatmaps for $Ca^{2+}$ responses in CaMKII+ (upper) and GAD67+ (bottom) neurons, before and after HNK injection in the same set of mice. Responses aligned to struggling onset.

(B) Quantification of onset to struggle onset (0 s) in $Ca^{2+}$ responses for CaMKII+ and GAD67+ neurons.

(C) Quantification of peak struggle onset (0 s) in $Ca^{2+}$ response for CaMKII+ and GAD67+ neurons.

(D) Identification of excitatory [*n* = 249 cells/10 mice] and inhibitory [*n* = 58 cells/8 mice] neurons *in vivo* in HNK-injected mice.

(E) sEPSC amplitude, area and decay time in c-Fos-positive E-Neurons.

(F) sEPSC area and decay time in fast-spiking GAD67-positive neurons (FSN).

(G) sEPSC area and decay time in regular-spiking GAD67-positive neurons (RSN).

Unpaired *t*-test (B - G).
